# From definition to discovery: metabolite markers of high temperature in green grape berries

**DOI:** 10.64898/2026.06.02.729726

**Authors:** Xi Lucie Zhan, Caroline Mauve, Fatma Lecourieux, Eric Gomès, Erwan Chavonet, Josep Valls Fonayet, Bertrand Gakière, Cyril Abadie, Pierre Pétriacq, David Lecourieux

## Abstract

Understanding how plants respond to high temperature is critical under global warming. Metabolite markers can provide insights into stress-responsive mechanisms and help guide strategies to maintain crop quality. However, heat-associated metabolite markers in grape berries remain poorly defined, particularly at the green stage, a critical phase of berry development during which early metabolic perturbations can influence subsequent ripening and ultimately determine berry composition and quality. Here, we applied berry-scale heat treatments of eight durations of two major wine cultivars, Cabernet Sauvignon and Merlot. Untargeted LC-MS profiling revealed both conserved and cultivar-dependent responses to heat. Based on these patterns, three time points were selected for targeted GC-MS analysis, and subsequent statistical analyses identified robust “cultivar-common heat markers”: glycine decreased, whereas galactinol increased consistently across time points and cultivars. “Cultivar-dependent heat markers” were identified: xylose, lyxose, citrulline, quinic acid, and glutamine, that consistently distinguished CS and Merlot fruits under heat stress. Notably, xylose, lyxose, citrulline, and quinic acid also differentiated the two cultivars under ambient conditions, underscoring their potential as stable cultivar-discriminating metabolites. Together, these results reveal dynamic metabolic remodeling in grape berries under heat stress, particularly in amino acid, nitrogen, central carbon metabolism, raffinose family oligosaccharides pathway and the glutathione-ascorbate cycle.

## INTRODUCTION

Anthropogenic greenhouse gas emissions are driving global warming, with global temperatures already ∼1.5 °C above pre-industrial levels and expected to continue rising (IPCC, 2023). This trend is accompanied by more frequent and intense heatwaves, with extreme temperatures above 40 °C increasingly recorded worldwide. Such events, exemplified by the so-called “Heat Dome”, have caused extensive plant damage and pose a growing threat to crop productivity and global food security (Kacprzyk et al. 2025).

Grapevine (*Vitis vinifera*) ranks as the third most economically significant crop worldwide and holds substantial socio-cultural value, underpinning the production of wine, table grapes, and raisins (Van Leeuwen et al. 2024). Beyond its economic significance, grapevine serves as a perennial model crop for scientific research: its obligate grafting to phylloxera-resistant rootstocks provides insights into graft union formation (Gautier et al. 2019), while clonal vegetative propagation offers opportunities for epigenetic studies (Fortes and Gallusci 2017). Moreover, grape berries are non-climacteric fruits, with ripening independent of ethylene regulation, making them an excellent model for investigating non-ethylene-mediated metabolic processes (Dai et al. 2016; Colombié et al. 2023). Extensive genomic resources further support their use for identifying molecular markers of environmental adaptation (This et al. 2006; Chen et al. 2025).

Plants have evolved complex mechanisms to cope with environmental fluctuations, often requiring reconfiguration of metabolic networks to maintain cellular homeostasis. Metabolites act as a bridge between genotype and phenotype, providing direct insights into gene functions and offering a powerful lens into biochemical and molecular mechanisms (Obata and Fernie 2012; Shen et al. 2023). More than 200,000 plant metabolites have been identified, broadly classified into primary and specialized metabolites. Primary metabolites sustain growth and development, whereas specialized metabolites contribute to metabolic diversity, defense, and adaptation, often in tissue- or stage-dependent ways (Wang et al. 2019; Shen et al. 2023). Importantly, both primary and specialized metabolites shape fruit organoleptic traits such as flavor, color, and aroma (Poni et al. 2018). This pleiotropic role underscores the importance of investigating rich metabolite networks under environmental stress.

Traditional abiotic stress studies often apply heat or drought to the whole-plant scale, but in vineyards, stress can also act locally at the cluster level. Sunlight exposure, canopy architecture, and viticultural practices such as leaf removal alter the berry microclimate, often elevating berry temperature and as a consequence, modifying diverse metabolic responses (Pieri and Fermaud 2005; Pieri et al. 2016). Organic acids, particularly malic acid, are highly sensitive to temperature. In Syrah berries, heat applied at *véraison* and during ripening accelerated malate degradation, whereas night-time or continuous day-and-night warming reduced diurnal variation and mitigated malate loss. These effects were not observed at the green stage, revealing that temperature-dependent malate contents depend on both developmental stage and diurnal cycles (Sweetman et al. 2014). A decrease in malic acid under high temperature was also observed in Cabernet Sauvignon berries exposed to berry-scale treatment, when heat was applied at *véraison* and ripening and metabolites were analyzed at harvest (Lecourieux et al., 2017). Other metabolites also respond strongly to heat, with increase in galactinol (Pillet et al. 2012) and decrease in anthocyanins (Mori et al. 2005, 2007; Lecourieux et al. 2017). The diversity of these metabolic responses highlights that berries activate multiple pathways under heat stress, raising the possibility that certain metabolites or metabolic classes could serve as reliable markers of heat perception or adaptation.

The concept of biomarkers, first defined in medicine as measurable indicators of biological processes, is considered in plants as quantifiable molecular indicators that can be associated with performance-and quality-related traits (Fernandez et al. 2016). In grapevine, metabolite markers are often assessed alongside physiological parameters to predict candidates linked to abiotic stress responses such as drought (Patin *et al*. 2025), to biotic stress resistance (Chavonet et al. 2025), or to agronomic traits such as successful graft union formation (Loupit et al. 2022). While effective for accessible traits, this approach is less applicable to berry-scale abiotic stress, for which no standardized physiological parameters or tools are available. Moreover, berry quality reflects the integration of multiple traits that cannot be captured by a single measure but are ultimately governed by metabolite composition (Poni et al. 2018). Stress-responsive metabolites can therefore serve as biomarkers, particularly when comparative analyses of cultivars with different heat sensitivities are undertaken, as single-cultivar measurements may not be sufficiently informative. Such comparisons enable the identification of both common and cultivar-dependent metabolites under stress, deepening insights into berry responses and support breeding strategies for stress-resilient cultivars.

The study by Lecourieux *et al*. (2017) showed that heat treatment at the green stage delayed *véraison* and altered berry composition at maturity, underscoring that heat exposure during early development can exert long-lasting effects on berry physiology and metabolism. In our previous work, heat exposure of ripening grape berries resulted in an “increase-then-decrease” pattern of polyphenols with increasing heat duration (Zhan et al. 2026). By contrast, heat applied at the green stage, when berries are undergoing cell division and expansion, did not trigger dynamic changes in polyphenols, although phenolic acids, flavanols, and flavonols were more abundant at this stage than at ripening. This suggests that other metabolites may play a more prominent role in mediating heat responses at the early developmental stage. Nevertheless, despite these insights, the effects of heat at the green stage have not yet received comprehensive attention, and their broader implications for berry quality and resilience remain underexplored.

In this context, the present study focuses on Cabernet Sauvignon and Merlot, two of the most widespread grape varieties worldwide. These two cultivars were selected for their putative different heat responses and assumed, distinct differences in ripeness and wine quality balance under climate warming (Van Leeuwen et al. 2019). Fruiting cuttings were grown in a greenhouse and subjected to berry-scale heat treatments (Lecourieux et al. 2017), generating ∼10 °C differences between control and heated berries at the green stage. Metabolic profiling was performed using a combination of untargeted LC-MS, targeted and untargeted GC-MS, integrating targeted metabolite quantitation with the broad coverage of untargeted analysis (Głuchowska et al. 2025). This approach enabled the characterization of a wide spectrum of primary and specialized metabolites, providing a panoramic snapshot of berry metabolite composition across control and heat treatments.

## MATERIALS AND METHODS

### Plant Material, heat treatment and sampling

Fruiting cuttings of *Vitis vinifera* L. cv Cabernet Sauvignon (clone #169) and Merlot (clone #343) were grown in a greenhouse under standardized conditions (Ollat et al., 1998). Plants were grown in 0.5 L pots filled with an equal mix of perlite, sand, and vermiculite, irrigated four times daily with nutrient solution, and maintained with 16 leaves and one cluster. Uniform plants were selected.

To simulate temperature increases seen in sun-exposed berries in vineyards, localized heat treatment (HT) was applied to grape clusters (Lecourieux et al. 2017). Plants were kept in a cooling greenhouse at ∼27 °C, while cluster temperature was raised by ∼10 °C during the day (08:00-16:00) for 10 days using radiators. Leaves and roots were insulated with polystyrene to restrict heating to clusters. Sampling followed the protocol described in Zhan et al. (2026)), using the same material as in the present study.

### Untargeted LC-MS metabolite extraction and processing

Approximately 10 mg (± 2%) of finely powdered, lyophilized material was aliquoted into 1.1 mL micronic tubes (MP32033L, Micronic, Lelystad, Netherlands) and randomly distributed across a 96-well micronic rack (MPW51001BC6, Micronic, Lelystad, Netherlands). Metabolite extraction was performed using an automated workflow developed at the MetaboHUB-Bordeaux Metabolome Facility. Briefly, samples underwent sequential extraction with two solvents: 300 µL of 80% ethanol supplemented with 0.1% formic acid (v/v), followed by 300 µL of 50% ethanol. The combined supernatants were subsequently filtered and preserved for downstream analyses, as previously described (Zhan et al. 2026). Quality control (QC) samples were generated by pooling equal-volume aliquots from all extracted supernatants. The pooled QC was subsequently aliquoted into micronic tubes and systematically injected at regular intervals (every ten study samples) throughout the analytical sequence in a randomized order to monitor data quality and instrument stability.

Raw untargeted LC-MS data were processed using MS-DIAL (version 4.90) (Tsugawa et al. 2015). Data were acquired over a retention time range of 0-18 min, with a mass range of 50-1500 Da for both MS1 and MS2 (Data Dependent, DDA) spectra. Peak detection parameters were set to 10 mDa for MS1 and 25 mDa for MS2, and isotope recognition allowed a maximum charge state of 2. Smoothing was performed using a linear weighted moving average method (over 4 scans), with a minimum of 5 scans per peak and a minimum peak height threshold of 10,000. Peak spotting employed a mass slice width of 0.05.

Metabolite annotation was performed using in-house and FragHUB MSP database files (Dablanc et al. 2024), and MS/MS spectral matching with the following parameters: retention time tolerance (100), MS1 tolerance (0.01), MS2 tolerance (0.05), and an identification score cutoff of 70. Retention time was not used for scoring or filtering. The relative abundance cutoff was set to 0, and only the top candidate per feature was retained. Alignment was performed using a QC file as reference. Parameters included retention time tolerance (0.1), MS1 tolerance (0.015), retention time factor (0.5), MS1 factor (0.5), and peak count filter (0) were set. Features were retained if detected in at least one group, with blank filtering applied (sample max/blank average ≥ 5; sample average/blank average ≥ 5). Gap filling was enabled. Isotope label tracking was disabled.

Normalization of aligned data was carried out using LOWESS combined with an internal standard (Methyl vanillate), with span = 0.7, minimum size = 0.11, and maximum size = 1. Final curation of LC-MS metabolome retained features with signal-to-noise ratio (S/N) > 10 and QC-based coefficient of variation (CV) < 30%.

### Targeted GC-MS metabolite extraction and analysis

Approximately 5 mg of freeze-dried, powdered tissue was transferred to 2-mL Safelock tubes and extracted in 1 mL of pre-cooled solvent (water:acetonitrile:isopropanol, 2:3:3, v/v/v) supplemented with ribitol (4 mg/L) as an internal standard (IS). Extractions were carried out for 10 min at 4 °C with agitation at 1500 rpm (Eppendorf Thermomixer). Insoluble residues were removed by centrifugation at 13,500 rpm for 10 min. 100µl of supernatants were evaporated at 35 °C for 4 h in a SpeedVac concentrator and stored at −70 °C until analysis. Blank tubes underwent the same protocol to serve as controls.

For chemical derivatization, 100 µL extracts were thawed, re-dried for 2 hours, and treated with 10 μL of methoxyamine hydrochloride (20 mg/mL in pyridine) for 90 min at 30 °C under constant shaking. Subsequently, 90 μL of N-methyl-N-trimethylsilyl-trifluoroacetamide (MSTFA; Regis Technologies, Cat. No. 1-270590-200) was added, and the mixture was incubated for 30 min at 37 °C. After cooling, 90 μL of each derivatized sample was transferred to Agilent vials. After 4 h, 1 μL aliquots were injected in splitless mode into an Agilent 7890B gas chromatograph coupled to an Agilent 5977A mass spectrometer, equipped with a DB-5MS column (30 m + 10 m Integra-Guard, Agilent, Ref. 122-5532G). Highly abundant metabolites were additionally measured with injections in split mode (ratio 1:30).

The oven was programmed at 60 °C for 1 min, increased at 10 °C/min to 325 °C, and held for 10 min. Helium was used as carrier gas at 1.1 mL/min constant flow. Temperatures were set at 250 °C for the injector, 290 °C for the transfer line, 230 °C for the ion source, and 150 °C for the quadrupole. Mass spectra were collected in electron-impact mode (50-500 m/z) after a solvent delay of 5.9 min. Sample order was randomized. External retention index calibration was performed using a fatty acid methyl ester (FAME) mixture (C8-C30).

Raw files were processed with AMDIS (http://chemdata.nist.gov/mass-spc/amdis/). Metabolite identification relied on the Agilent Fiehn GC/MS Metabolomics RTL Library (June 2008), which contains electron-impact spectra and retention indices for ∼700 reference compounds. Peak integration was performed in MassHunter Quantitative Analysis (Agilent) under both splitless and split modes; automated integration was manually inspected and corrected when necessary. Resulting peak areas were compiled into a single dataset, normalized to ribitol and sample dry weight, and expressed as relative abundances.

### Untargeted GC-MS analysis for the tentative identification of new markers

GC-MS data were processed using MS-DIAL (version 4.90) following the same workflow applied for the untargeted LC-MS analysis described above. Putative metabolite identification was performed by comparing mass spectra and retention indices with entries in the Agilent Fiehn GC/MS Metabolomics RTL Library described above.

### Statistical analysis

Following IS normalization, untargeted LC-MS data were square-root transformed and Pareto scaled, and normality was assessed using MetaboAnalyst 5.0 (Pang et al. 2022). Targeted GC-MS data were evaluated for normality using the Shapiro-Wilk test in R (v4.5.2) for each metabolite within each condition, with 88-99% of metabolites normally distributed (*p* > 0.05). Similarly, 73.6-97.5% of untargeted GC-MS fragments followed normal distributions, supporting parametric analyses. Statistical analyses were performed in R. At each time point, differences between heat-treated and control groups were tested using unpaired *t*-tests, with a fold-change threshold of 1.5. Two-way ANOVA assessed treatment and genotype effects, and metabolites showing significant genotype × treatment interactions were further analyzed by post-hoc comparisons. Significance was set at *p* < 0.05 after false discovery rate (FDR) correction, while uncorrected p-values were retained for exploratory analysis of untargeted GC-MS data. Pathway analysis was conducted using the KEGG database for *Arabidopsis thaliana* in MetaboAnalyst 5.0, with manual validation of metabolite annotations and enrichment analysis performed in R. Multivariate analyses and visualizations were carried out in R: heatmaps (ComplexHeatmap, circlize packages), volcano plots and correlation analyses (ggplot2), PLS-DA (ropls), and Venn diagrams (eulerr).

## RESULTS

### 1. Time-resolved metabolomic responses to high temperature

To assess temporal metabolic responses, grapevine berries were subjected to localized heat treatment (ht) of varying durations: short-term exposures (30min, 1h, 2h, 4h, and 1d) and long-term exposures (2d, 5d, and 10d). At each time point, comparisons between control (ctl) and HT-treated berries were performed for both CS and Merlot (Figure 1A, B).

**Figure 1.**
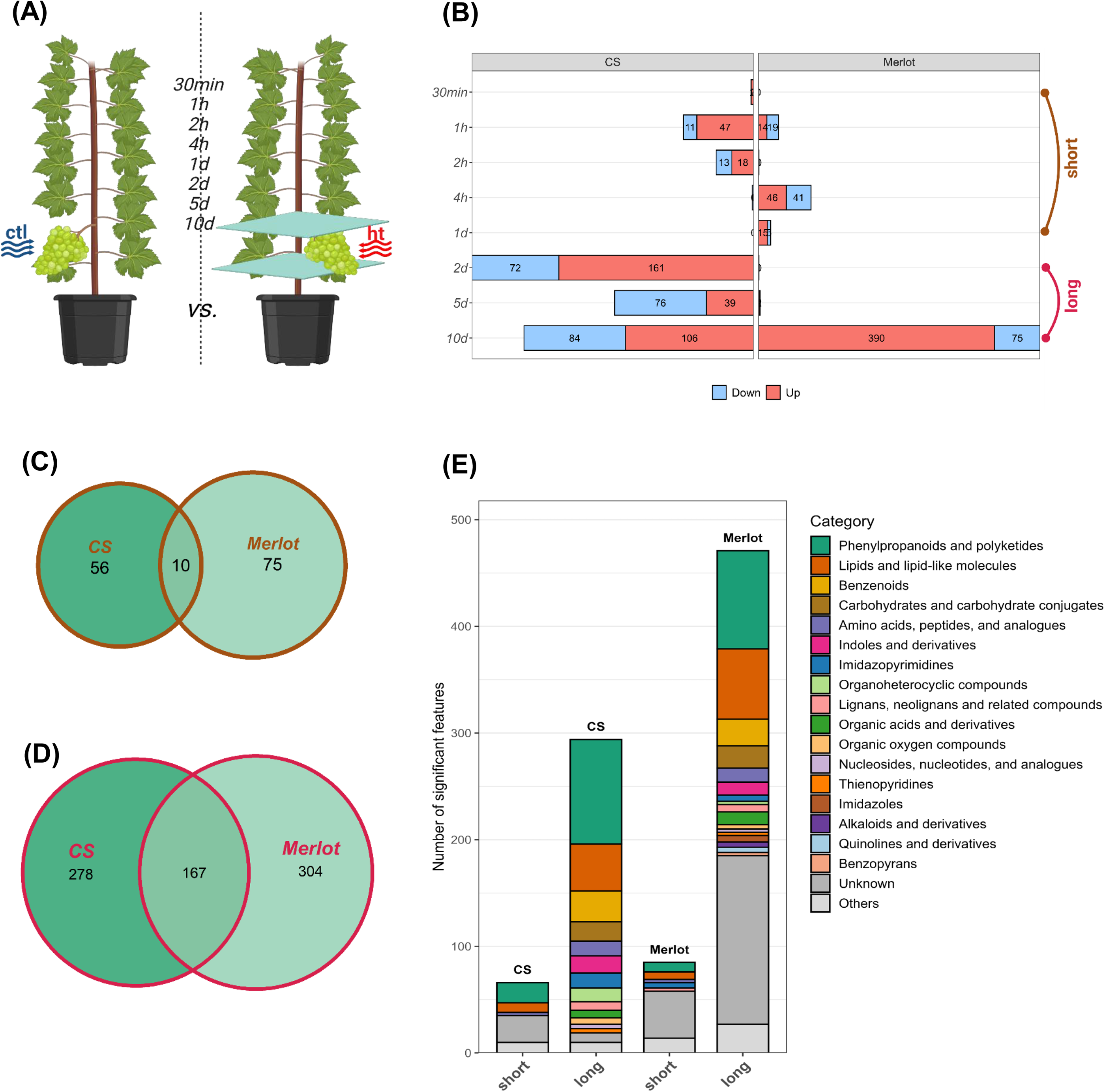
**(A)** Schematic representation of localized heat treatment in herbaceous grape berries. **(B)** Number of significant features from untargeted analysis at each time point (ht/ctl) in CS and Merlot. Features were considered significant with *p* < 0.05 (FDR) and fold change (ht/ctl) > 1.5. **(C)** Venn diagrams of significant features under short-term and **(D)** long-term treatments in CS and Merlot. **(E)** Categories of significant features in both cultivars under short- and long-term treatments. Abbreviations: ht, heat treatment; ctl, control; CS, Cabernet-Sauvignon.

Statistical analyses were performed on 5843 filtered features (from an initial 15038 detected features). Differentially accumulated features (DAFs) were defined as features identified by untargeted LC-MS that met both statistical and biological significance thresholds: a two-sided Student’s *t*-test with FDR-adjusted *p* < 0.05 and an absolute fold change (ht/ctl) > 1.5 (Figure 1B). In CS, short-term HT resulted in 92 DAFs, with 67 increased and 25 decreased. In Merlot, 141 DAFs were detected, of which 76 increased and 65 decreased. The largest number of DAFs was observed at 2d in CS and at 10d in Merlot (Figure 1B). With prolonged exposure, responses intensified: CS displayed 538 DAFs (306 increased, 232 decreased), whereas Merlot exhibited 469 DAFs (393 increased, 76 decreased).

Comparison of the two cultivars revealed limited overlap under short-term HT, with only 10 shared DAFs, including members of the pyran and amino acid classes, as well as tricoumaroyl spermidine (Figure 1C; Supplementary Table 1). In contrast, long-term HT resulted in 167 shared DAFs, predominantly from the benzenoid, lignan/neolignan, and lipid classes (Figure 1D; Supplementary Table 1). Unique DAFs specific to each cultivar and treatment duration are provided in Supplementary Table 1.

Classification of DAFs by metabolite category revealed distinct time-dependent responses (Figure 1E). In CS, short-term HT primarily affected phenylpropanoids and polyketides, lipids and lipid-like molecules, and amino acids or related derivatives. With long-term HT, the response broadened to include benzenoids, carbohydrates, indoles, imidazopyrimidines, and other organoheterocyclic compounds. In Merlot, short-term HT influenced similar categories, phenylpropanoids and polyketides, lipids, and amino acids or related derivatives, but also includes imidazopyrimidines and lignans/neolignans. Long-term HT further expanded these responses to encompass benzenoids, carbohydrates, indoles, organoheterocyclic compounds, and additional lignan/neolignan derivatives, alongside the categories already altered during short-term exposure.

### 2. Cultivar-dependent metabolomic profiles under high temperature

To investigate cultivar-dependent responses to HT, we compared CS and Merlot at each time point (Figure 2A). The number of significant features varied across treatments, ranging from as few as 6 at 1d to as many as 480 at 1h. Among short-term treatments, the largest responses were detected at 1h (480 features) and 4h (255 features), whereas under long-term HT, the strongest response was observed at 5d (448 features) (Figure 2B). Based on these results, three representative time points, 1h, 4h, and 5d were selected for pathway enrichment analysis.

**Figure 2.**
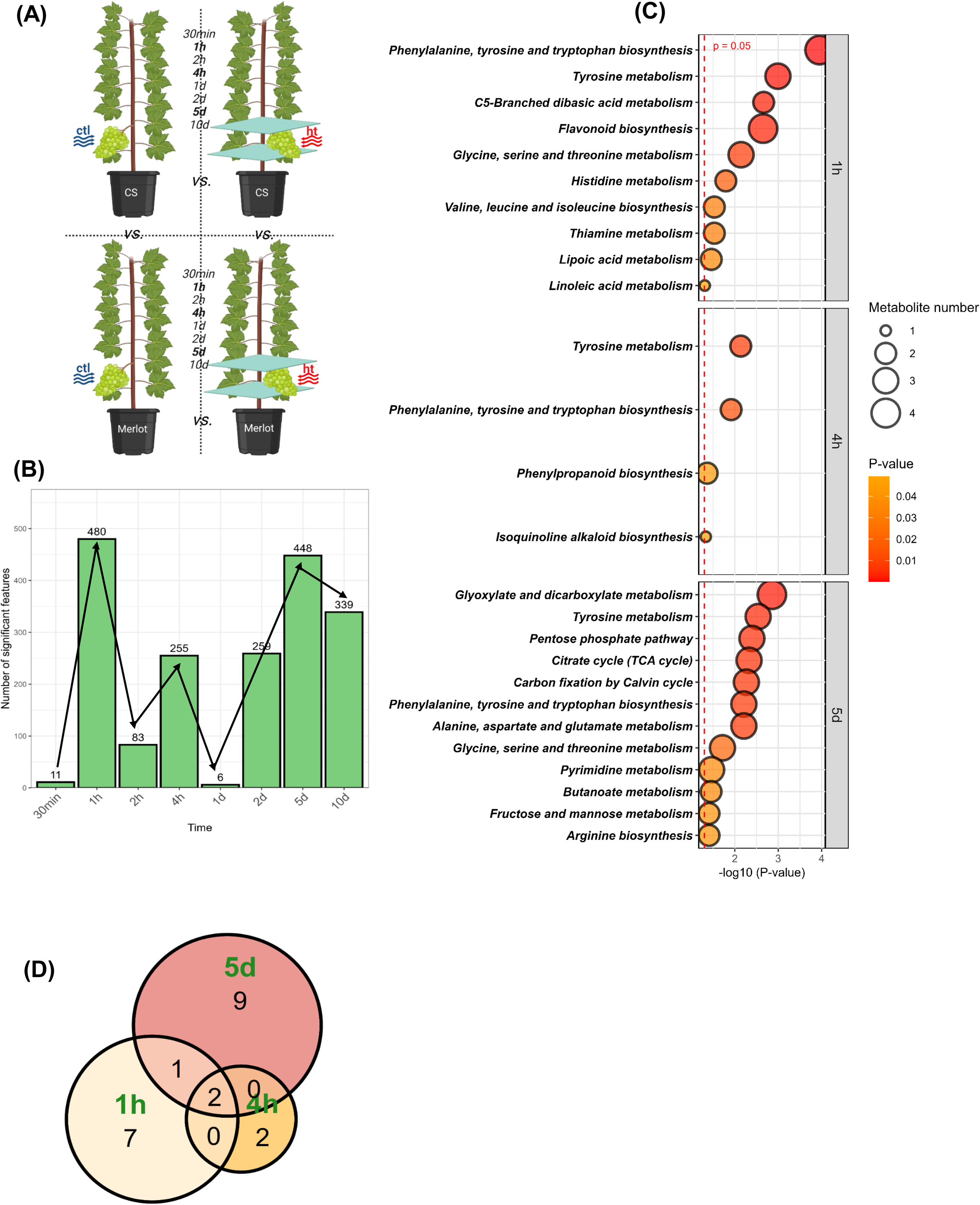
**(A)** Schematic representation of localized heat treatment in herbaceous grape berries of two cultivars. **(B)** Differential features from untargeted LC-MS analysis at each time point identified by two-way ANOVA (cultivar × treatment), with FDR-adjusted *p* < 0.05 considered significant. **(C)** Pathway enrichment analysis showing metabolic pathways involved at selected time points, with *p* < 0.05 considered significant. **(D)** Venn diagrams of enriched pathways at three selected time points.

Pathway enrichment revealed distinct temporal patterns (Figure 2C). At 1h, features mapped to 10 pathways, mainly involving amino acid metabolism (e.g., phenylalanine, tyrosine and tryptophan biosynthesis, tyrosine), with additional pathways related to secondary metabolism (e.g., flavonoid) and lipid metabolism (e.g., lipoic and linoleic acids). At 4h, four pathways were enriched, including amino acid metabolism and phenylpropanoid biosynthesis. At 5d, 12 pathways were identified, spanning central carbon metabolism (e.g., TCA cycle, pentose phosphate pathway, Calvin cycle), amino acid metabolism, and carbohydrate metabolism (e.g., fructose and mannose). Across the three selected time points, two pathways were consistently enriched: phenylalanine, tyrosine and tryptophan biosynthesis, and tyrosine metabolism (Figure 2C, D). Overall, the two-way ANOVA revealed that CS and Merlot exhibit distinct metabolic trajectories under heat stress, with consistent differences in amino acid-related pathways, suggesting that regulation of aromatic amino acid metabolism plays a central role in cultivar-dependent heat responses.

### 3. Cultivar-common heat markers

To obtain targeted quantitation of metabolic markers associated with HT in grape berries, targeted GC-MS analysis was performed at selected time points (1h, 4h, and 5d) in both cultivars, yielding 81 confidently annotated metabolites (Supplementary Table 2).

In CS (Figure 3A), 8 metabolites changed significantly after 1h of HT, with 7 increased (mainly amino acids such as tyrosine and threonine) and 1 decreased (glycine). At 4h, 11 metabolites were altered, including increases in sugars (e.g., sucrose, galactinol) and decreases in organic acids and phosphorylated intermediates (e.g., malic acid, G6P, F6P). After 5d, 14 metabolites were significantly changed, with broad increases across sugars, organic acids, and amino acids, while glycine again decreased. In Merlot (Figure 3B), 3 metabolites changed significantly after 1h of HT, with 2 increased (aspartic acid and 3-phosphoglycerate) and 1 decreased (glycine). At 4h, 13 metabolites were altered, with increases in sugars and sugar derivatives (e.g., galactinol, glucono-δ-lactone, gluconic acid, galactonic acid) and decreases in amino acids and sugar phosphates (e.g., glycine, aspartic acid, glutamic acid, G6P, F6P, 3-phosphoglycerate). After 5d, 18 metabolites were significantly changed, characterized by broad increases in amino acids (e.g., glutamine, threonine, isoleucine, histidine, arginine), sugars (e.g., raffinose, galactinol), and organic acids, while a smaller set of metabolites, including glycine and sugar phosphates, decreased.

**Figure 3.**
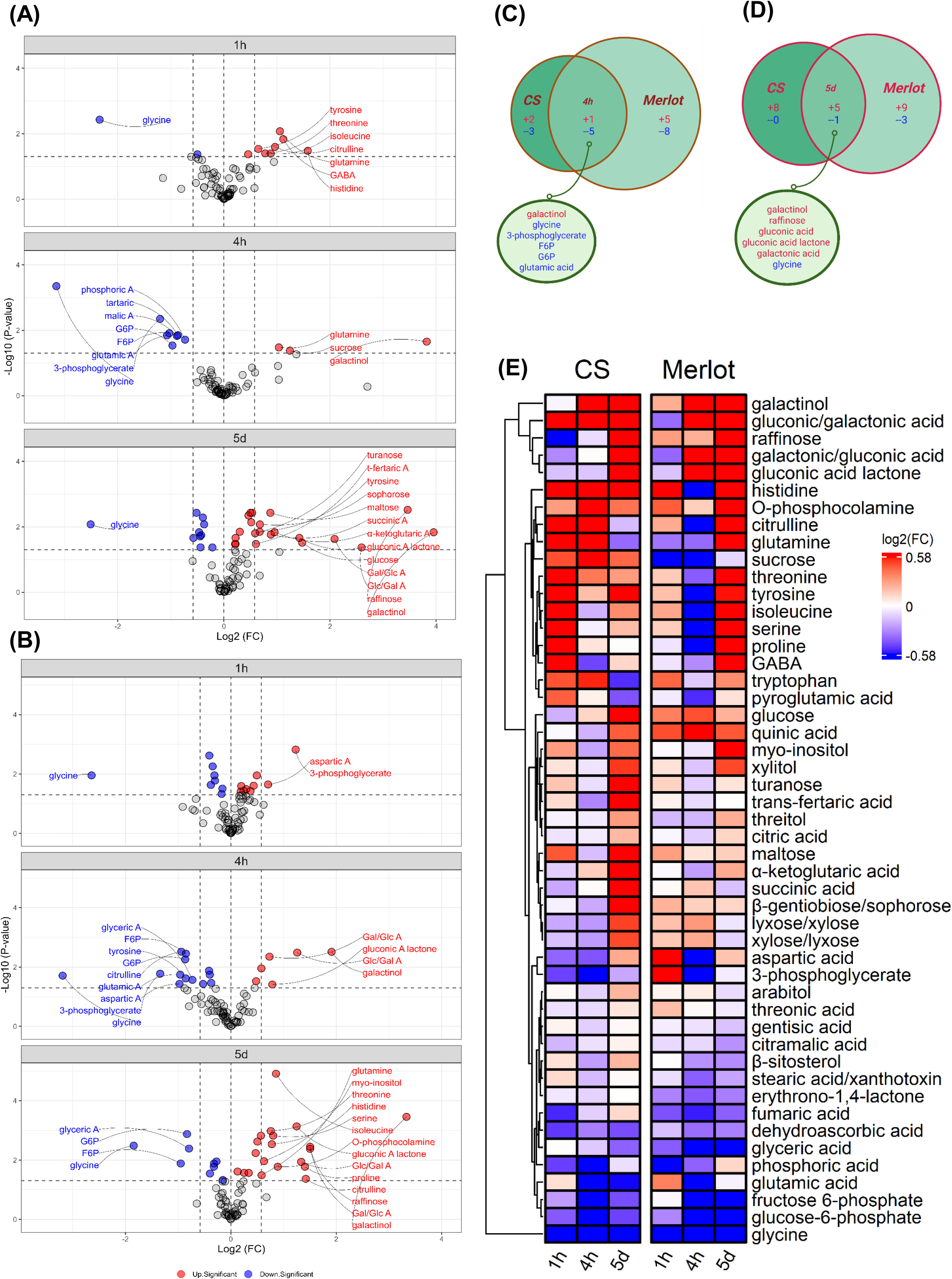
Volcano plots showing significant metabolites from targeted GC-MS analysis at three time points (1h, 4h, and 5d) based on *t*-test analysis (ht/ctl), with *p* < 0.05 (FDR correction) considered significant, in CS **(A)** and Merlot **(B).** Only metabolites with *p* < 0.05 (FDR) and fold change |FC| > 1.5 are labeled in the volcano plots. Venn diagrams of significant metabolites (*p* < 0.05, FDR; |FC| > 1.5) at 4h **(C)** and 5d **(D)** in CS and Merlot. **(E)** Heatmap of selected metabolites with *p* < 0.05 (FDR) and FC > 1.5 at least at one time point in CS or Merlot. **Abbreviations:** A, acid; G6P, glucose-6-phosphate; F6P, fructose-6-phosphate; Gal, galactonic; Glc, gluconic; t, trans.

To highlight common metabolite markers across cultivars, we focused on metabolites showing consistent and concordant changes in both CS and Merlot at the same time points (Figures 3C, D). After 1h of HT, glycine was the only metabolite consistently decreased in both cultivars (Figure 3A, B). At 4h, galactinol increased, while five metabolites, glycine, 3-phosphoglycerate, F6P, G6P, and glutamic acid, consistently decreased. After 5d of HT, five metabolites, galactinol, raffinose, glucono-δ-lactone, gluconic acid, and galactonic acid increased in both cultivars, whereas glycine again decreased. Metabolites levels significantly altered (*p* < 0.05, FDR, |FC| > 1.5) in at least one cultivar or time point were further examined across all time points and cultivars (Figure 3E), providing a broader view of their evolutions. Galactinol and raffinose accumulated rapidly in Merlot, with increases already evident at 1h of HT, whereas in CS these changes appeared later. In contrast, amino acids began to increase from 1h in CS but showed a weaker response in Merlot, where several decreased at 4h before rising more strongly by 5d. Ultimately, the magnitude of amino acid accumulation was greater in Merlot than in CS. Several metabolites clustered together that belong to carbohydrate metabolism (glucose, maltose, turanose, β-gentiobiose/sophorose, lyxose/xylose) and sugar alcohols (xylitol, threitol), along with central carbon intermediates of the TCA cycle (citric acid, α-ketoglutaric acid, succinic acid) and quinic acid, trans-fertaric acid, and myo-inositol were clustered. Overall, these metabolites showed relatively modest changes under short-term HT but exhibited a marked increase after 5d in CS, whereas the increase was less evident in Merlot. A distinct cluster of metabolites showed a decreasing trend in both CS and Merlot. These included TCA cycle intermediates (e.g., fumaric acid), antioxidant compounds (e.g., dehydroascorbic acid), sugar phosphates (e.g., fructose-6-phosphate, glucose-6-phosphate, glyceric acid), inorganic phosphates (e.g., phosphoric acid), and amino acids (e.g., glutamic acid, glycine). The decline was already evident under short-term HT and remained pronounced at 5d.

### 4. Cultivar-dependent heat markers

In addition to heat-increased and heat-decreased markers shared across cultivars, we identified a third category of metabolites whose responses to high temperature differed significantly between CS and Merlot, termed “cultivar-dependent heat markers”. To identify these, two-way ANOVA was performed on the targeted GC-MS dataset at three time points (Figure 4A). A total of 17 significant metabolites were identified at 1h, 9 metabolites at 4h, and 28 metabolites at 5d (Figure 4A; Supplementary Table 2).

**Figure 4.**
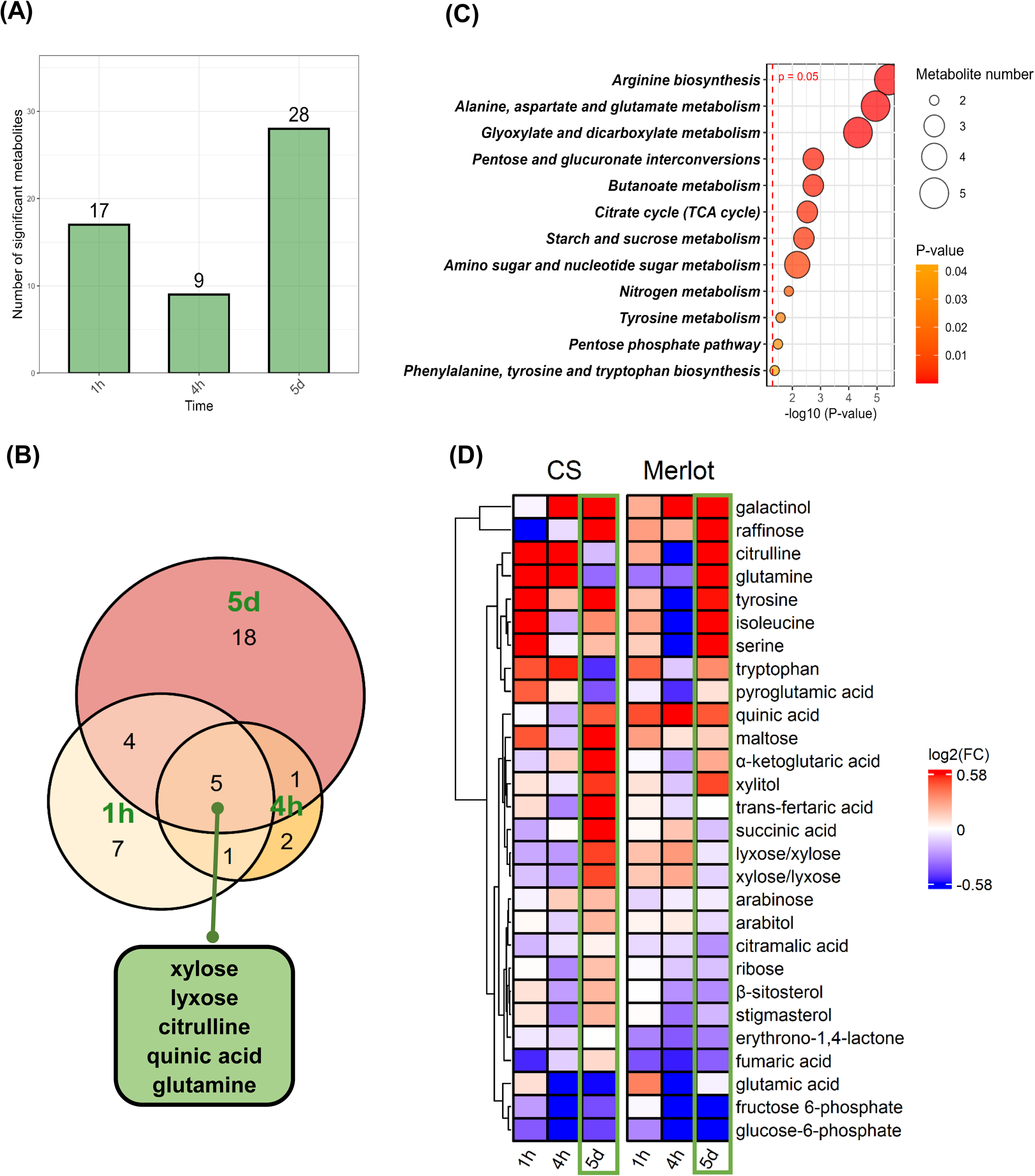
**(A)** Number of significant metabolites from targeted GC-MS analysis based on two-way ANOVA (cultivar × treatment) at three time points (1h, 4h, and 5d), with *p* < 0.05 (FDR) considered significant. **(B)** Venn diagram of significant metabolites from two-way ANOVA across the three time points. **(C)** Enrichment pathway analysis showing significant pathways involved at 5d (*p* < 0.05). **(D)** Heatmap of 28 significant metabolites (identified by two-way ANOVA at 5d), with their fold changes (log2 fold change) at 1h and 4h also shown.

Overlap analysis across time points revealed five metabolites consistently detected at all three stages (Figure 4B): xylose, lyxose, citrulline, quinic acid, and glutamine. Pathway enrichment of the 5d metabolite set indicated significant involvement of multiple processes, particularly amino acids (e.g., arginine, alanine, tyrosine pathways), carbohydrate (e.g., TCA cycle, pentose phosphate pathway, starch and sucrose metabolism), as well as butanoate and nitrogen metabolisms (Figure 4C).

Several of the overlapping metabolites displayed contrasting cultivar-dependent patterns at 5d (Figure 4D). Xylose and lyxose increased in CS but decreased in Merlot. Citrulline and glutamine showed opposite trends in two cultivars, decreased in CS but increased in Merlot. Quinic acid increased in both cultivars, though to different extents. In addition to previously described temporal patterns in metabolites (Figure 3E), ribose, β-sitosterol, and stigmasterol also exhibited contrasting behaviors at 5d, with increases in CS and decreases in Merlot (Figure 4D).

### 5. Cultivar-dependent heat markers in control conditions

To further explore whether cultivar-dependent heat markers also differ under baseline conditions, we evaluated their behavior in control berries of CS and Merlot. PLS-DA was applied to assess separation between cultivars and to identify key metabolites contributing to discrimination across time points (Figure 5A).

**Figure 5.**
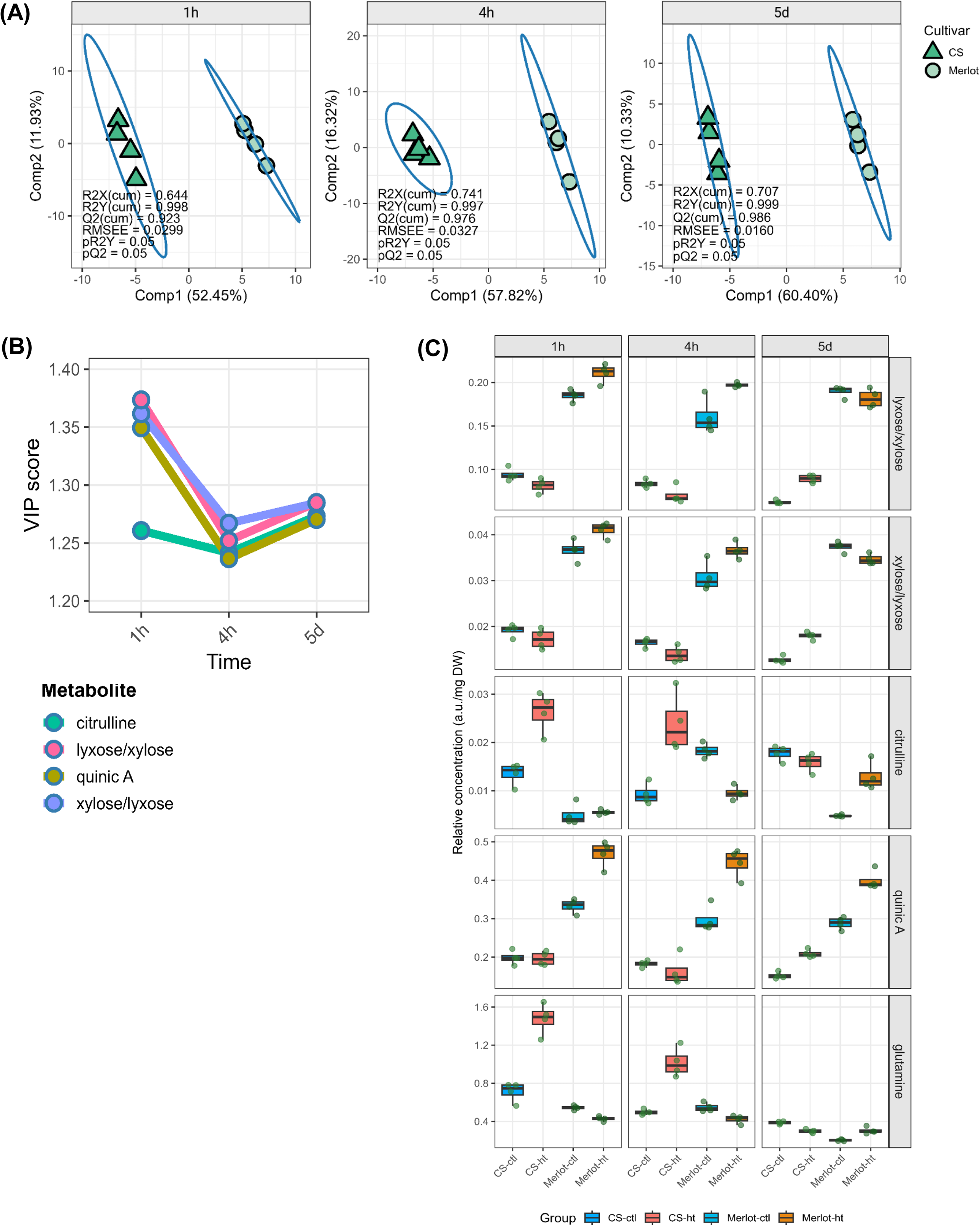
**(A)** PLS-DA score plots of (targeted GC-MS) of CS and Merlot under control conditions at three time points: 1 hour (1h), 4 hours (4h), and 5 days (5d). Model performance is indicated by R²X, R²Y, Q², RMSEE, pR²Y, and pQ² values. **(B)** VIP trajectories of four selected metabolite markers across the three time points. **(C)** Relative concentrations (normalized to ribitol and dry weight, DW) of five metabolite markers. Abbreviations: A, acid.

At each time point, samples clustered distinctly according to cultivars (Figure 5A), with separation primarily along the first component (Comp1), which explained the majority of the variance (52.45%, 57.82%, and 60.40% at 1h, 4h, and 5d, respectively). Model quality parameters indicated robust performance, with high cumulative R²Y values (0.998-0.999) and strong predictive power (Q² = 0.923-0.986). Cross-validation showed low RMSEE values (0.0160-0.0327), and permutation tests confirmed model validity (pR²Y = 0.05, pQ² = 0.05).

The Variable Importance in Projection (VIP) analysis from the PLS-DA model identified metabolites with VIP > 1 that contributed most strongly to cultivar separation across the three time points (Figure S1). At 1h, several carbohydrate-related metabolites (e.g., arabinose, maltose, sophorose) and amino acids (e.g., tryptophan, tyrosine) showed high discriminatory power. At 4h, both carbohydrate metabolites (e.g., sucrose, maltose, turanose) and amino acids (e.g., tryptophan, citrulline) remained strong contributors, together with shikimic acid. By 5d, the separation was more pronounced, with consistent contributions from amino acids (e.g., tryptophan, citrulline) and sugars (e.g., maltose, sophorose, xylose/lyxose). Notably, tryptophan was among the top contributors at all three time points, underscoring its consistent role in cultivar discrimination. Overall, metabolites associated with amino acid and carbohydrate metabolism were the most important drivers of separation, though their relative contributions varied with time.

Five metabolites previously identified as key markers of cultivar-dependent heat responses, citrulline, xylose, lyxose, quinic acid, and glutamine were further evaluated within the PLS-DA framework (Figures 4B and 5B). Four of these (citrulline, lyxose, xylose, and quinic acid) consistently showed VIP > 1 across all time points (Figure 5B). Comparison of their relative concentrations revealed that citrulline, lyxose, xylose, and quinic acid also differed significantly between CS and Merlot under control conditions (Figure 5C, Supplementary Table 3), whereas glutamine responses under HT appeared independent of baseline variation.

### 6. Integrative correlation analysis between targeted cultivar-common metabolite markers and untargeted LC-MS features

We selected two metabolite markers consistently responsive to HT in both cultivars: galactinol (a heat-increased marker) and glycine (a heat-decreased marker). Using these markers as reference metabolites, we explored untargeted metabolomics data by Pearson correlation to identify additional features potentially associated with their behaviors (Figure S2).

Galactinol showed significant negative correlations with several specialized metabolites-like features (e.g., tricoumaroyl spermidine, acacetin, bilobalide) and positive correlations with phenolics and related glycosides (e.g., sinapyl alcohol, syringin, kaempferol derivatives) (Figure S2A). Glycine, in turn, was positively correlated with multiple organic acids and conjugates, including amino acid derivatives and spermidines, while showing a negative correlation with sinapyl alcohol (Figure S2B). In CS, galactinol was positively correlated with flavonoid glycosides (e.g., genistein and phenylethyl glucoside) but negatively correlated with a wide range of metabolites, including amino acids, alkaloids, and fatty acid derivatives (Figure S2C). Glycine exhibited broad positive correlations with phenolics, amino acid derivatives, and glucosinolates, while showing negative correlations with organic acids and lipid derivatives (e.g., phosphatidylethanolamine alkeryl, fatty acyl hexoside) (Figure S2D).

### 7. Tentative exploratory of new markers based on untargeted GC-MS analysis

To explore novel heat markers, an untargeted GC-MS analysis was conducted, leading to the detection of 992 fragments. Cultivar-common fragments were screened by comparing control and HT at three selected time points in both CS and Merlot. In CS, a total of 667 differentially accumulated fragments (DAFs) were identified, among which 578 DAFs showed decreased pattern under HT. In Merlot, 333 DAFs were detected, with 246 exhibiting decreased levels following HT (Figure S3A). Across the three time points, 25 and 33 DAFs were consistently detected in CS and Merlot (Figure S3B, S4), respectively. Notably, four DAFs were common to both cultivars at all time points (Figure S3B). The temporal profiles of these common fragments (annotated as 817, 819, 162 and 313) revealed a consistent decrease in abundance across all HT durations and were selected for further tentative identification (Figure S3C, S4).

Cultivar-dependent fragments were identified using two-way ANOVA (genotype × condition) across three selected time points in both CS and Merlot. A total of 95, 108 and 120 DAFs were detected at 1h, 4h and 5d, respectively (Figure S5A). Among these, 11 DAFs exhibited a significant genotype × condition interaction consistently across all time points (Figure S5B). Their temporal profiles revealed contrasting heat response patterns between CS and Merlot (Figure S5C). These 11 DAFs were selected for further tentative identification.

Compound identification was performed by comparing mass spectra and retention characteristics with spectral libraries and reference data. N-ethylglycine was tentatively assigned as a cultivar-common marker, whereas dihydroxybenzoic acid was identified as a cultivar-dependent marker (Table 1; Supplementary Table 4). N-ethylglycine exhibited a consistent decrease across all three time points in both CS and Merlot. In contrast, dihydroxybenzoic acid displayed divergent response patterns between cultivars, showing an “increase-then-decrease” trend in CS and a “decrease-then-increase” pattern in Merlot under HT (Figure 6).

**Figure 6.**
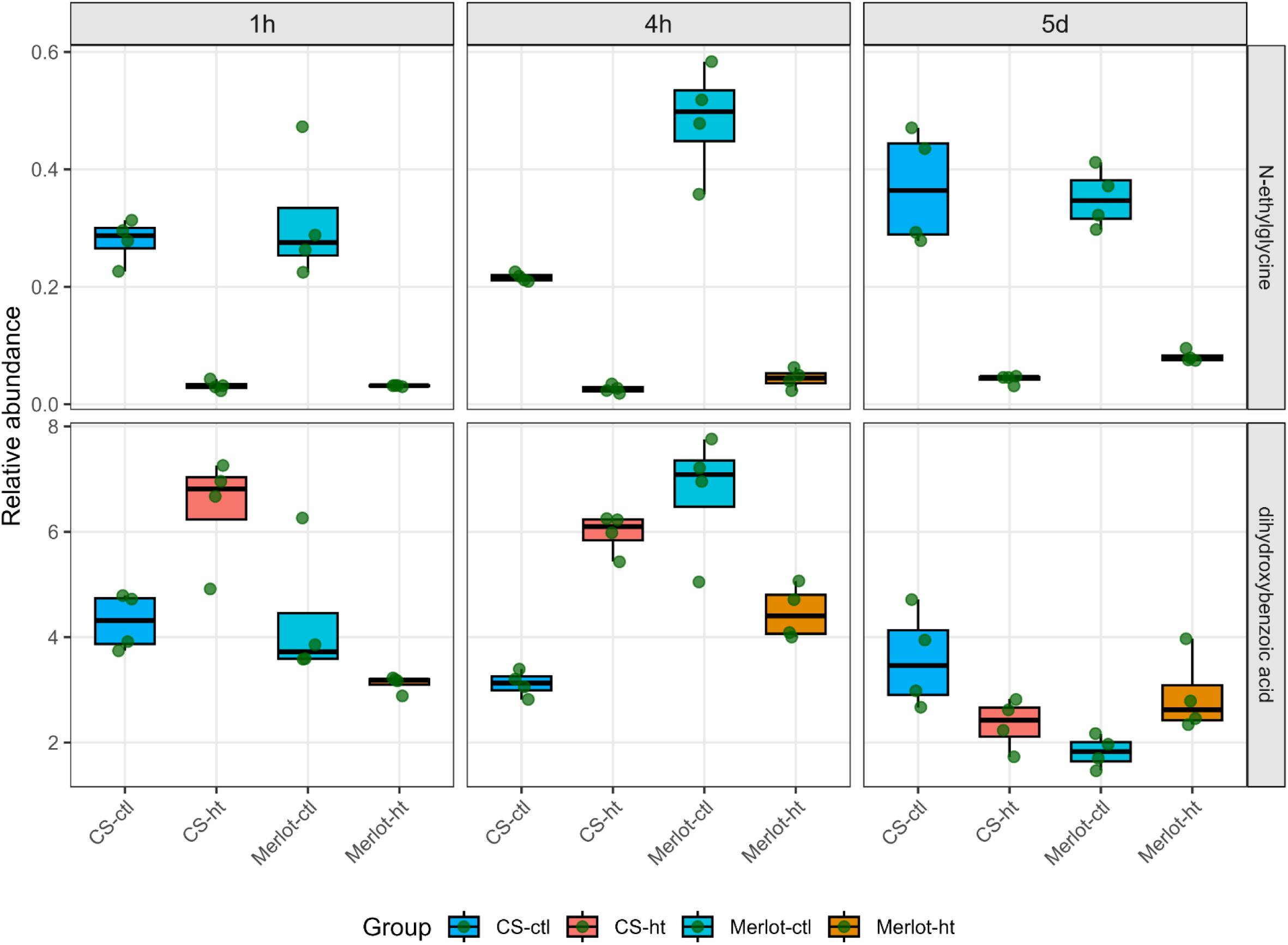
Ribitol-normalized relative abundances of N-ethylglycine and gentisic acid in Cabernet-Sauvignon (CS) and Merlot under control (ctl) and heat treatment (ht) conditions at 1h, 4h, and 5d.

**Table 1.**
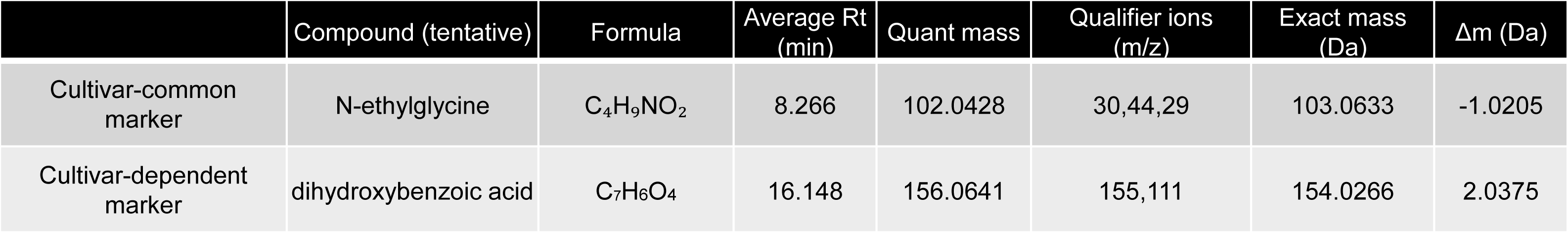
Identification of discriminant metabolites in GC-MS untargeted analysis. Tentatively identified metabolites associated with cultivar-common and cultivar-dependent markers are listed along with their chemical formulas and mass spectrometric characteristics. Quantification was based on ribitol-normalized peak areas. Compound identification was achieved by comparison of mass spectra and retention behavior with spectral libraries and reference data. Reported parameters include qualifier ions (used for confirmation), exact mass, and mass deviation (Δm) expressed in Daltons.

## DISCUSSION

### 1. Heat duration-dependent metabolomic profiles

Our time-resolved metabolomics analysis revealed distinct, duration-dependent metabolic responses to HT in CS and Merlot (Figure 1E), with phenylpropanoids and polyketides among the most affected classes (Figure 1). In our previous targeted profiling of polyphenols at the green stage, covering phenolic acids, flavanols, and flavonols, no significant changes were observed (Zhan et al. 2026). In contrast, the present untargeted LC-MS approach detected additional heat-responsive metabolites, notably flavonoid C- and O-glycosides (Supplementary Table 1). This discrepancy reflects methodological coverage: targeted LC-MS focuses on a defined set of aglycones (e.g., quercetin, kaempferol), whereas untargeted analyses capture a broader spectrum of structurally diverse, decorated flavonoids typically absent from targeted panels.

Flavonoid glycosides were markedly altered under HT, suggesting that glycosylation contributes to metabolic remodeling in response to HT. Glycosylation enhances flavonoid solubility, stability, and bioactivity, thereby supporting plant adaptation to abiotic and biotic stress (Zhao et al. 2024; Dong et al. 2025). This process is catalyzed by UDP-glycosyltransferases (UGTs), many of which are stress-inducible. Functional studies in other species support this role: in rice, UGT706F1, upregulated by MYB61, enhances flavonoid glycoside accumulation and thermotolerance (Zhao et al. 2025), while the BZIP16-UGT2 module contributes to drought and salt tolerance via enhanced ROS scavenging. In tea (*Camellia sinensis*), the heat-responsive *Cs*UGT73A17 catalyzes the formation of 7-O-glucosides, with both transcript levels and enzymatic activity increasing under heat (Su et al. 2018). In grapevine, *Vv*UGT125 is highly expressed in berries and upregulated in shoot tips under drought (Hou et al. 2025), and transcriptomic analyses identified heat-responsive UGTs in green berries, including *Vv*UGT (VIT_05s0062g00340), which was induced after one day of HT in CS (Lecourieux et al. 2017). Together, the transcript-level evidence from grapevine and functional studies in other crops, combined with our metabolomic detection of glycosylated flavonoids, supports a role for UGT-mediated glycosylation in grape berry response to high temperature.

Metabolomic profiles revealed that lipids and lipid-like molecules were strongly affected by HT in both CS and Merlot under short- and long-term treatments, mainly involving non-free fatty acid classes such as fatty acyl glycosides, glycerophosphoethanolamines, glycerophosphoserines, and terpenoid derivatives. This pattern suggests broad membrane-associated adjustments to heat stress (Figure 1D, Supplementary Table 1). The role of lipids in thermotolerance is well established in plants (Zheng et al. 2011; Li et al. 2015; Narayanan et al. 2016). Plants mitigate heat stress through multiple strategies, including affecting the membranes by changing the fluidity and permeability (Saidi et al. 2010). During heat priming in *Arabidopsis thaliana*, levels of major phospholipids such as glycerol-3-phosphate, glycerophosphorylcholine, and glycerophosphoethanolamine increase, alongside upregulation of lipid-turnover enzymes including *LCAT4* (AT1G45201, encoding a lecithin-cholesterol acyltransferase-like enzyme with phospholipase A1 activity) and AT4G29070 (encoding a phospholipase A2-type enzyme) (Serrano et al. 2019). This coordinated response suggests enhanced phospholipid recycling that supports membrane stability under heat stress. Such remodeling may also contribute to thermomemory and is consistent with interactions between glycerophospholipid, terpenoid, and tocopherol pathways, promoting the synthesis of protective compounds such as glycine betaine and vitamin E (Porfirova et al. 2002; Ashraf and Foolad 2007; Serrano et al. 2019). In grapevine, transcriptomic data (Lecourieux et al. 2017) showed that a phosphoesterase family protein (VIT_06s0009g03350) is significantly upregulated in green berries after prolonged heat exposure. Together, these metabolomic and transcriptomic observations suggest that phospholipid turnover is dynamic under heat stress, with phosphoesterase family proteins likely contributing to the recycling of phospholipid intermediates and the maintenance of membrane lipid homeostasis, thereby supporting heat response in grape berries.

Compared to short-term HT, long-term HT induced broader metabolic diversification (Figure 1D), with benzenoids significantly affected only under prolonged treatment in both cultivars. Benzenoids are key plant volatiles (Lv et al. 2024), in grapes and wines, volatile benzenoids such as 2-phenylethanol (rose/floral), benzaldehyde (almond/nutty), eugenol (clove/spice), and vanillin (vanilla/woody) are well documented. However, because samples were freeze-dried, most volatiles were likely lost during preparation. Accordingly, the benzenoid-like features detected by untargeted LC-MS likely represent non-volatile intermediates or precursors rather than aroma compounds (Collins et al. 2015, 2015; Slaghenaufi et al. 2019), explaining why many annotations fall within broad benzenoid classes without matching known grape aroma compounds (Supplementary Table 1). Nevertheless, benzenoids are reported as stress-responsive metabolites in other species, such as *Achillea millefolium* under heat stress (Liu et al. 2021a), suggesting that benzenoid metabolism, including volatile derivatives, may contribute to grape berry heat responses and potentially influence aroma.

Indoles and their derivatives were predominantly affected by long-term HT (Figure 1D). Derived from tryptophan metabolism, indole-3-acetic acid (IAA) is a central auxin regulating plant development (Radwanski and Last 1995). In grape berries, auxin controls ripening: exogenous IAA delays ripening by inhibiting cell wall degradation, while endogenous IAA declines after anthesis and its conjugation is required for ripening initiation (Davies & Robinson 1997; Lin et al. 2025; Bottcher, Keyzers, Boss & Davies 2010; Cawthon & Morris 1982). Heat stress has been linked to IAA-mediated growth regulation (Li et al. 2021; Abdullah et al. 2023; Bianchimano et al. 2023). Consistently, transcriptomic data in grapevine show that HT represses IAA-amido synthetase (VIT_07s0005g00090) and induces IAA-amino acid hydrolases (VIT_11s0016g02700, VIT_08s0007g02740), suggesting that heat disrupts IAA conjugation and delays onset of ripening as a consequence (Lecourieux et al. 2017). Our metabolomic evidence of altered indole metabolism supports this mechanism.

### 2. Cultivar-dependent metabolomic responses under HT

As previously reported, CS responds more rapidly than Merlot to heat at the ripening stage. Consistently, our metabolomic analysis at the green stage showed that CS displayed more significant features under both short- and long-term HT (Figure 1B), indicating cultivar-dependent heat responses. Two-way ANOVA (cultivar × treatment) revealed that short-term effects were most pronounced at 1h and 4h, while long-term effects peaked at 5d (Figure 2B). Pathways including phenylalanine, tyrosine, and tryptophan biosynthesis, together with phenylpropanoid metabolism, were significantly affected at all time points, highlighting the sensitivity of primary metabolism and cultivar-dependent responses (Figure 2C). At 1h, differences were mainly associated with amino acid metabolism (aromatic and branched-chain) and lipid pathways, suggesting distinct strategies in nitrogen remobilization and membrane adjustment. At 4h, fewer metabolic pathways differentiated the cultivars, indicating a transient convergence. By 5d, differences expanded to central carbon and carbohydrate metabolism (e.g., TCA cycle, pentose phosphate, glyoxylate/dicarboxylate, fructose/mannose), along with multiple amino acid pathways and butanoate metabolism, reflecting long-term remodeling of energy supply and carbon-nitrogen balance. Overall, CS and Merlot differ not only in speed but also in the breadth of their metabolic responses to HT, with early nitrogen and lipid adjustments followed by broader carbon-energy remodeling, supporting the identification of cultivar-specific metabolic markers.

### 3. Cultivar-common heat markers

Glycine emerged as a consistent heat-decreased marker, showing significant reductions in both CS and Merlot at all time points, in agreement with vineyard observations across other cultivars (Wang et al. 2025). This supports its role as a central metabolic node under HT (Figure 3, 7). In plants, glycine is a key intermediate of serine-glycine one-carbon metabolism within photorespiration, contributing to nucleotide synthesis and methylation. In mitochondria, two glycine molecules are converted into one serine via the glycine decarboxylase complex (GDC) and serine hydroxymethyltransferase (SHMT), while SHMT also catalyzes the reverse reaction through one-carbon transfer to tetrahydrofolate (Bauwe et al. 2010) (Figure 7). Both enzymes are abundant in the mitochondrial matrix of green tissues, whereas GDC activity is negligible in non-green tissues such as potato tubers (Mouillon et al. 1999). The GDC-SHMT system catalyzes the rapid degradation of glycine exported from peroxisomes during photorespiration (Douce et al. 2001). In grape berries, transcriptomic analyses identified five DEGs encoding *Vv*SHMT isoforms (VIT_00s0211g00160, VIT_00s0211g00090, VIT_00s0211g00120, VIT_00s0211g00070, VIT_00s0211g00100) (Lecourieux et al. 2017).

**Figure 7.**
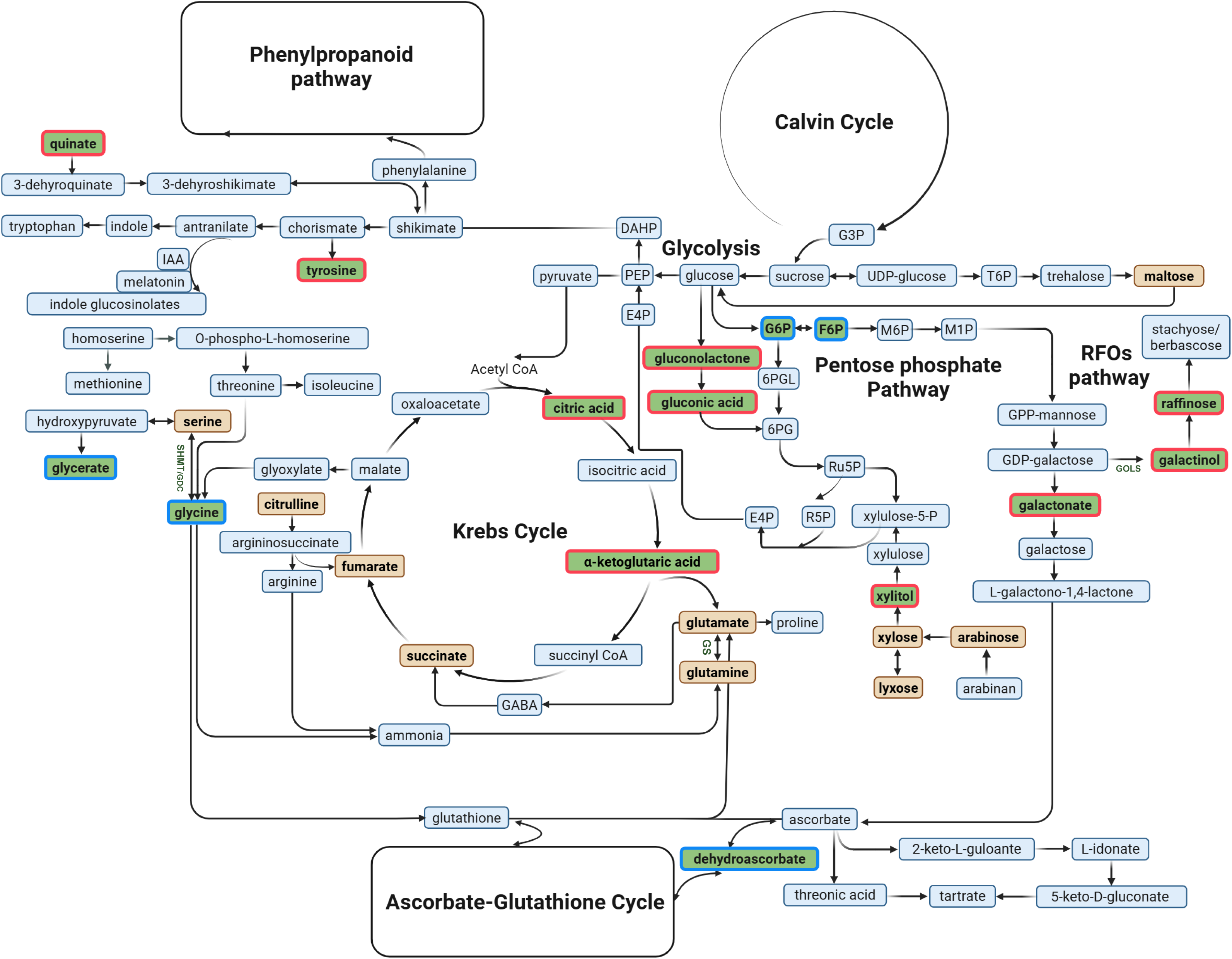
Metabolic biosynthesis pathways illustrating both common and cultivar-specific heat-responsive pathways in CS and Merlot. The list of common significant metabolites is from targeted analysis after 5 days of treatment (see Figure 3D), with additional significant metabolites included (*p* < 0.05, FDR, but |FC| < 1.5). **Box colors:** green = significant metabolites based on *t*-test (ht/ctl); green with red edges = significantly accumulated after 5 days of treatment; green with blue edges = significantly decreased after 5 days of treatment; orange = significant metabolites identified by two-way ANOVA when comparing CS and Merlot after 5 day of treatment (see Figure 4). **Abbreviations**: IAA, indole-3-acetic acid; GABA, γ-aminobutyric acid; DAHP, 3-deoxy-D-arabino-heptulosonate 7-phosphate; PEP, phosphoenolpyruvate; E4P, erythrose-4-phosphate; G3P, glyceraldehyde-3-phosphate; T6P, trehalose-6-phosphate; G6P, glucose-6-phosphate; F6P, fructose 6-phosphate; 6PGL, 6-phosphoglucono-δ-lactone; 6PG, 6-phosphogluconate; Ru5P, ribulose-5-phosphate; R5P, ribose-5-phosphate; Xu5P, xylulose-5-phosphate.

Interestingly, these *Vv*SHMT genes were largely repressed in green berries after 1d, 7d, and 14d of HT, with three isoforms consistently repressed across all treatments. Reduced SHMT expression, together with decreased glycine levels, suggests a constraint in serine-glycine interconversion, potentially limiting substrate availability for GDC. This may restrict photorespiratory flux and one-carbon metabolism, with downstream effects on growth, stress responses, and amino acid homeostasis.

Galactinol is identified as a robust heat-increased marker in green berries. It increased after 4h and 5d of HT in both CS and Merlot (Figure 3). This agrees with Pillet *et al*. (2012), who showed that *Vv*GOLS1, encoding a galactinol synthase (Figure 7), is strongly heat-induced and transactivated by *Vv*HsfA2. While that study focused on *véraison* and ripening stages in CS, our results extended this response to the green stage and across cultivars, confirming galactinol as a consistent heat-induced marker. Raffinose, another raffinose family oligosaccharides (RFO), accumulated significantly after 5d of HT in both cultivars (Figure 3D, Figure 7). Consistently, Lecourieux *et al*. (2017) reported two raffinose synthase genes (VIT_17s0000g08960 and VIT_14s0066g00810), among which VIT_17s0000g08960 was consistently induced in green berries after 1d, 7d, and 14d of HT, with transcript levels increasing as heat duration was prolonged. This time-dependent regulation likely explains why significant raffinose accumulation was only observed at 5d in our data. In other plants, galactinol and raffinose function as cellular osmoprotectants, scavenging hydroxyl radicals and thereby protecting against oxidative damage under abiotic stress (Nishizawa et al. 2008; Yan et al. 2022). In tomato, galactinol also promotes cold tolerance by promoting jasmonic acid (JA) biosynthesis. It facilitates the conversion of linolenic acid, an unsaturated fatty acid important for membrane stability, into JA, thereby reinforcing JA-mediated signaling under stress (Hu et al. 2013; Liu et al. 2024). Together, our results identify galactinol as a reliable marker of heat response and highlight the central role of the RFO pathway in metabolic and signaling adaptation to heat stress in grape berries.

### 4. Cultivar-dependent heat markers

To identify metabolite markers distinguishing cultivars, CS and Merlot were compared using two-way ANOVA at each HT time point. The greatest differences occurred under very short-term (1h) and long-term (5d) HT (Figure 4A). Across all time points, five metabolites, xylose, lyxose, citrulline, quinic acid, and glutamine were consistently identified as cultivar-dependent markers (Figure 4B).

Xylose and lyxose exhibited contrasting patterns after 1h, 4h, and 5d of HT in the two cultivars (Figure 4D). In CS, their levels decreased initially and then increased, whereas in Merlot, they increased first and subsequently declined. Xylose, a pentose sugar, is the main building block of xylan, the predominant hemicellulose in plant cell walls. With its β-1,4-linked backbone and side-chain decorations, xylan reinforces cell wall strength and flexibility through interactions with cellulose and lignin, making xylose essential for proper wall structure buildup and plant growth (York and Oneill 2008). An increase in xylose content was observed in *Scrophularia striata* subjected to drought stress (Falahi et al. 2018). Similarly, xylose accumulation has been suggested as a tolerance mechanism against drought stress, contributing to osmotic adjustment and homeostasis in canola shoots (Viana et al. 2022). In contrast, xylose content decreased in coffee leaves under heat stress (Lima et al. 2013). (Du et al. 2011) reported that xylose increased in both *bermudagrass* and *Kentucky bluegrass* under HT. In our study, both sugars were more abundant in Merlot under control conditions (Figure 4C) and displayed opposite responses in two cultivars under HT, supporting their role as robust cultivar-dependent markers. These patterns suggest distinct cell wall remodeling and osmotic adjustment strategies between CS and Merlot. Lyxose has been detected in sweet potato but not in potato or cassava and could be associated with cell wall metabolism. Its coordinated behavior with xylose suggests a similar role in cultivar-dependent metabolic regulation (Figure 7) (Mutanda et al. 2026).

Citrulline emerged as a cultivar-dependent marker. This nonessential amino acid, abundant in watermelon (Citrullus vulgaris Schrad), functions as an effective osmoprotectant and antioxidant under abiotic stress (Song et al. 2020; Malambane et al. 2023). In desert watermelon (*Citrullus lanatus*), citrulline is constitutively high and contributes to osmotic balance and oxidative stress protection, while salinity stress instead promotes the production of other compatible solutes such as γ-aminobutyric acid (GABA), proline, and glutamine (Kusvuran et al. 2013). In contrast, melon (*Cucumis melo*) increases citrulline levels under salinity and drought, with greater accumulation in tolerant genotypes (Kusvuran et al. 2013). The divergence was also observed in our study, where citrulline accumulation differed between CS and Merlot across HT durations (Figure 5C), suggesting distinct cultivar-specific strategies for osmotic adjustment and redox regulation, although the underlying mechanisms remain unclear.

Quinic acid was identified as the fourth cultivar-dependent heat (Figure 4B). Its role in abiotic stress is not well defined. In eggplant, exogenous quinic acid enhanced resistance to biotic stress in a susceptible cultivar (Liu et al. 2022), while in tobacco cell cultures, quinic acid derivatives were induced by defense-priming agents (Mhlongo et al. 2014). Derivatives such as caffeoylquinic acids correlate strongly with antioxidant capacity and accumulate under heat stress in tomato leaves via *HsfB1* regulation (Paupière et al. 2020; Islam et al. 2024). Quinic acid derivatives have also been linked to antioxidant activity through their potential role as ligands of enzymes such as superoxide dismutase in *Carissa spinarum* (Liu et al. 2021b). In our study, the baseline level of quinic acid was higher in Merlot and increased largely under HT in Merlot but not in CS (Figure 5C). We speculate that the higher basal abundance of quinic acid in Merlot may facilitate its more rapid and pronounced induction under HT. While evidence for the direct role of quinic acid in abiotic stress response remains limited, our findings point to its potential contribution to cultivar-dependent heat responses in grape berries.

Glutamine was also identified as a cultivar-dependent heat marker. As a central amino acid in nitrogen metabolism, glutamine is regarded as promising markers for determining abiotic stress such as drought and salinity stress in plants (Yin et al. 2022; Kim-Teng Lee et al. 2023). Its biosynthesis *via* glutamine synthetase (GS) is critical for nitrogen assimilation through the glutamine synthetase-glutamate synthase (GS-GOGAT) cycle and supports stress tolerance in plants (Yin et al. 2022) (Figure 7). Gene expression analysis in contrasting durum wheat genotypes showed that the most tolerant genotype had higher GS activity and increased GS1 and GS2 expression under drought (Yousfi et al. 2016). Similarly, a proteomic study showed that chloroplastic GS2 was upregulated in a drought-tolerant wheat cultivar compared with a sensitive one (Cheng et al. 2016). These findings align with our metabolomic data, where glutamine increased under HT in CS but not in Merlot, suggesting cultivar-dependent GS regulation. In contrast, *Vv*GS2 (VIT_05s0020g02480) was repressed in CS green berries under HT (Lecourieux et al. 2017). The increase in glutamine despite transcript repression likely reflects the weak correlation between mRNA levels and metabolite abundance, as GS activity is often regulated post-transcriptionally (Yin et al. 2022). Another possible explanation for glutamine accumulation in CS under HT is reduced turnover within the GS/GOGAT cycle. Although GS transcripts were downregulated, GOGAT, which consumes glutamine to regenerate glutamate, was also strongly repressed in heated berries (VIT_12s0055g00620, VIT_15s0024g01030, VIT_16s0098g00290) (Lecourieux et al. 2017). In this case, the greater limitation on glutamine utilization likely outweighs reduced synthesis, indicating a shift in metabolic flux toward glutamine retention. Alternatively, heat stress promotes proteolysis (Callis 1995; Fan and Jespersen 2025), may increase the release of free amino acids such as glutamine, while reducing glutamine utilization for protein synthesis (Barnabás et al. 2008), further limiting its consumption, together favoring net accumulation. These processes may explain the contrasting glutamine responses of CS and Merlot, suggesting cultivar-dependent regulation of nitrogen flux under heat stress.

Except for glutamine, the other four cultivar-dependent heat markers, lyxose, xylose, quinic acid, and citrulline, showed predominant contributions in separating CS and Merlot under control conditions (Figure 5A,B), with VIP scores >1 at all three time points. This indicates that these metabolites are not only robust markers under HT but also differentiate the two cultivars in the absence of stress, thereby underscoring their potential as genetic markers for breeding strategies.

### 5. Integrative metabolomic approaches reveal complementary heat-responsive metabolic networks across LC-MS and GC-MS approaches

Here, we integrated LC-MS and GC-MS to exploit their complementarity. Galactinol and glycine emerged as cultivar-common heat markers and central hubs in correlation networks (Figure S2). Tricoumaroyl spermidine and glutathione were negatively correlated with galactinol but positively correlated with glycine (Figure S2A, S2B), indicating coordinated decreases under HT. Tricoumaroyl spermidine has been reported as an abundant phenolamide in reproductive organs such as tea flowers, pear, and rose pollen (Yang et al. 2012; Qiao et al. 2023; Chen et al. 2018), and phenolamides can exhibit higher antioxidant activity than flavonoids in bee pollen berries (Zhang et al. 2020). This suggests that glycine- and galactinol-centered networks may function as redox hubs under heat stress in grape berries. Glycine, a direct precursor of glutathione (a key antioxidant in redox homeostasis) (Noctor et al. 2024; Mullineaux and Rausch 2005), decreased under heat stress and was positively correlated with glutathione, implying that decreased glycine may constrain antioxidant capacity in heat-treated berries. In addition, bilobalide-like and silibinin-like (silybin-like) features showed similar correlation patterns (Figure S2C, S2D). Given their reported antioxidant properties, these compounds may contribute to redox regulation under heat stress (Kim et al. 2024; Wadhwa et al. 2022).

In addition, untargeted GC-MS tentatively identified N-ethylglycine as a cultivar-common marker, whereas dihydroxybenzoic acid appeared as a cultivar-dependent heat marker (Table 1). N-ethylglycine has been mainly reported in human studies, where it is associated with metastatic bone disease (Tsuruta et al. 2007). In plants, its occurrence and function remain poorly documented, with limited evidence such as its decrease in broad bean (*Vicia faba* L.) roots following nanoparticle exposure (Tian et al. 2020). In grapevine, Sun et al. (2018) showed that N-ethylglycine levels in Merlot leaves under frost stress were influenced by root temperature, decreasing under warm roots and increasing under cold roots. Biochemically, N-ethylglycine is a structural derivative of glycine. In animal systems, it is known as a metabolite of lidocaine that modulates glycinergic signaling by inhibiting glycine transporters, thereby increasing extracellular glycine levels (Werdehausen et al. 2015). Although this mechanism is not directly transferable to plants, it highlights a close relationship between N-ethylglycine and glycine metabolism. In our data, both compounds showed similar decreasing patterns under HT, supporting the hypothesis that N-ethylglycine reflects glycine pool dynamics rather than an independent metabolic pathway. However, due to the lack of established biosynthetic pathways in plants, further targeted validation is required to confirm its identity and clarify its role under stress conditions.

Dihydroxybenzoic acid (DHBA) was tentatively identified as a cultivar-dependent marker (Table 1). DHBA comprises several structural isomers, including 2,3-, 2,4-, 2,5-, 2,6-, and 3,4-dihydroxybenzoic acids. Among these, 2,5-DHBA (gentisic acid) was detected in the targeted GC-MS dataset but was not identified as a significant marker (Supplementary Table 2), suggesting that the DHBA signal in the untargeted dataset likely corresponds to other isomers. Although the roles of DHBA isomers in abiotic stress remain unclear, benzoic acid derivatives are closely linked to salicylic acid (SA)-related metabolism and plant defense. In tea (*Camellia sinensis*), the UDP-glycosyltransferase *Cs*UGT74B5 glycosylates benzoic acid derivatives, including 2,6-DHBA, a hydroxylated SA derivative, in a structure-dependent manner favoring ortho-dihydroxylated compounds (Lu et al. 2024). This glycosylation reduces levels of active free forms and is associated with decreased disease resistance, highlighting the regulatory role of DHBA conjugation in defense response. Similarly, in *Arabidopsis thaliana*, DHBA glycosylation modulates SA-associated immunity (Huang et al. 2018). In addition, DHBA compounds possess intrinsic redox activity, for instance, 2,3-DHBA exhibits antioxidant properties in chemical systems (Okabe and Kyoyama 2001), suggesting a potential contribution to redox processes in plant tissues.

Altogether, even without physiological or multi-omics data, the integration of untargeted LC-MS, targeted and untargeted GC-MS, combined with robust statistical analyses, enabled comprehensive characterization of metabolic remodeling and identification of potential heat-responsive markers in grape berries. These results demonstrate that metabolomics alone can serve as a powerful and autonomous approach to dissect plant responses to abiotic stress.

This study provides the first comprehensive, time-resolved metabolomic dissection of green grape berries under localized high temperature, uncovering both cultivar-common and cultivar-dependent metabolite markers. We identified glycine as a reliable heat-decreased marker and galactinol as heat-increased marker, highlighting photorespiration and the RFO pathway as conserved components of the fruit heat response. In parallel, five metabolites, xylose, lyxose, quinic acid, citrulline, and glutamine emerged as cultivar-dependent markers, offering new insights into the differential metabolic strategies employed by Cabernet Sauvignon and Merlot. Strikingly, four metabolites not only responded to heat but also differentiated the cultivars under control conditions, underscoring their potential as genetic markers for breeding. Taken together, our findings refine the concept of metabolite markers from simple descriptors of stress to dynamic indicators of metabolic remodeling and cultivar identity. In this study, we focused on seven metabolites that consistently and significantly responded to heat across all treatment durations, establishing them as robust metabolite markers. However, many other metabolites displayed significant changes only at one or two time points. Although excluded from our marker definition, these transient responses likely capture signaling events or reflect condition-dependent metabolic remodeling, involving processes such as turnover, transport, or rerouting of fluxes that become prominent under high temperature. Their temporal specificity makes them equally valuable for understanding the dynamics of heat stress responses, and future studies should investigate these metabolites to unravel the layered regulation of berry metabolism under high temperatures.

## ACKNOWLEDGEMENT

Xi Lucie Zhan was supported by a grant from the China Scholarship Council (CSC) and by the Agence Nationale de la Recherche (ANR) through the project PARASOL (ANR-20-CE21-0003). This research also received funding from the Grand Programme de Recherche Bordeaux Plant Sciences (GPR-BPS) and from the Bordeaux Metabolome Facility and the MetaboHUB (ANR-11-INBS-0010). The authors gratefully acknowledge Jean-Pierre Petit for assistance with fruiting cutting production, greenhouse experiments, and technical training. We also thank the Embassy of France in Beijing for supporting scientific exchanges at the Plateforme Métabolisme-Métabolome, Université Paris-Saclay. Special thanks are extended to Françoise Gilard and Florence Guérard for their assistance with platform training.

