## Supplemental figures for "From definition to discovery: metabolite markers of high temperature in green grape berries"

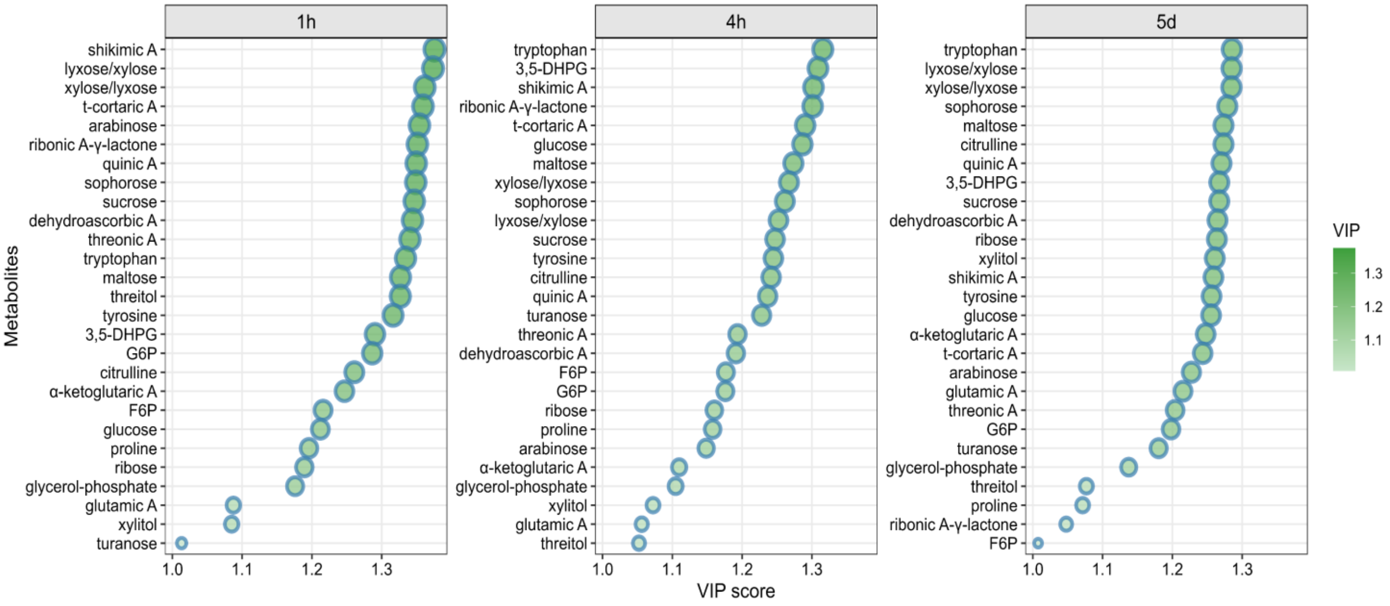


**Figure S1**. Variable importance in projection (VIP) scores of metabolites identified by targeted GC-MS at three time points under control condition. Metabolites with higher VIP scores contributed more strongly to the separation between cultivars. Results are shown for 1h, 4h, and 5d. The size and color intensity of the circles indicate the magnitude of the VIP score.

| | 4h (short-term) | 5d (long-term) |
|---|---|---|
| galactinol | Merlot | CS |
| glycine | Merlot | CS |

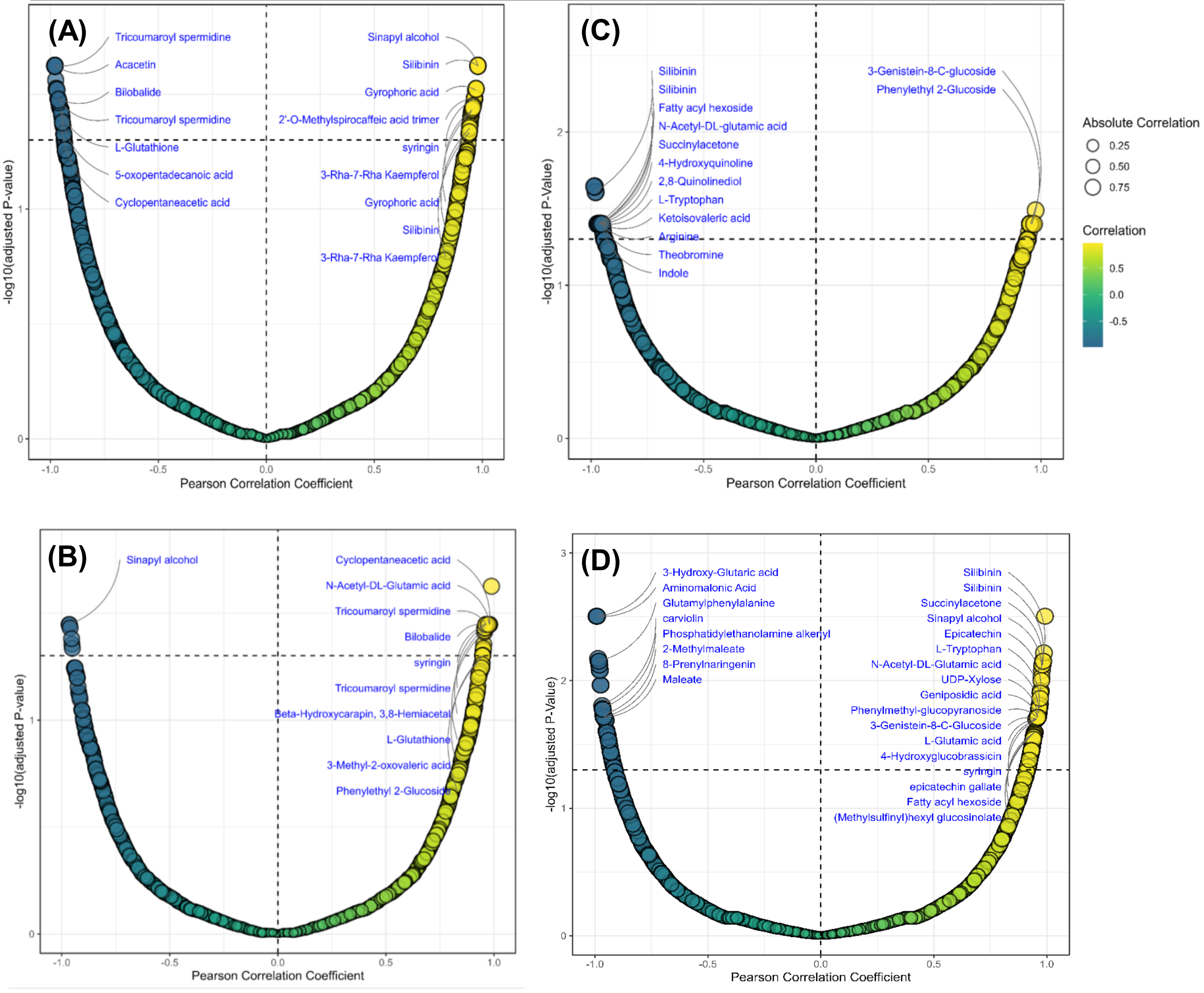


**Figure S2.** Pearson correlation coefficient analysis between **galactinol** and **glycine** (identified from targeted GC-MS) and features from the untargeted LC-MS dataset at both 4h and 5d. Correlations with $P < 0.05$ (FDR-adjusted) were considered significant.

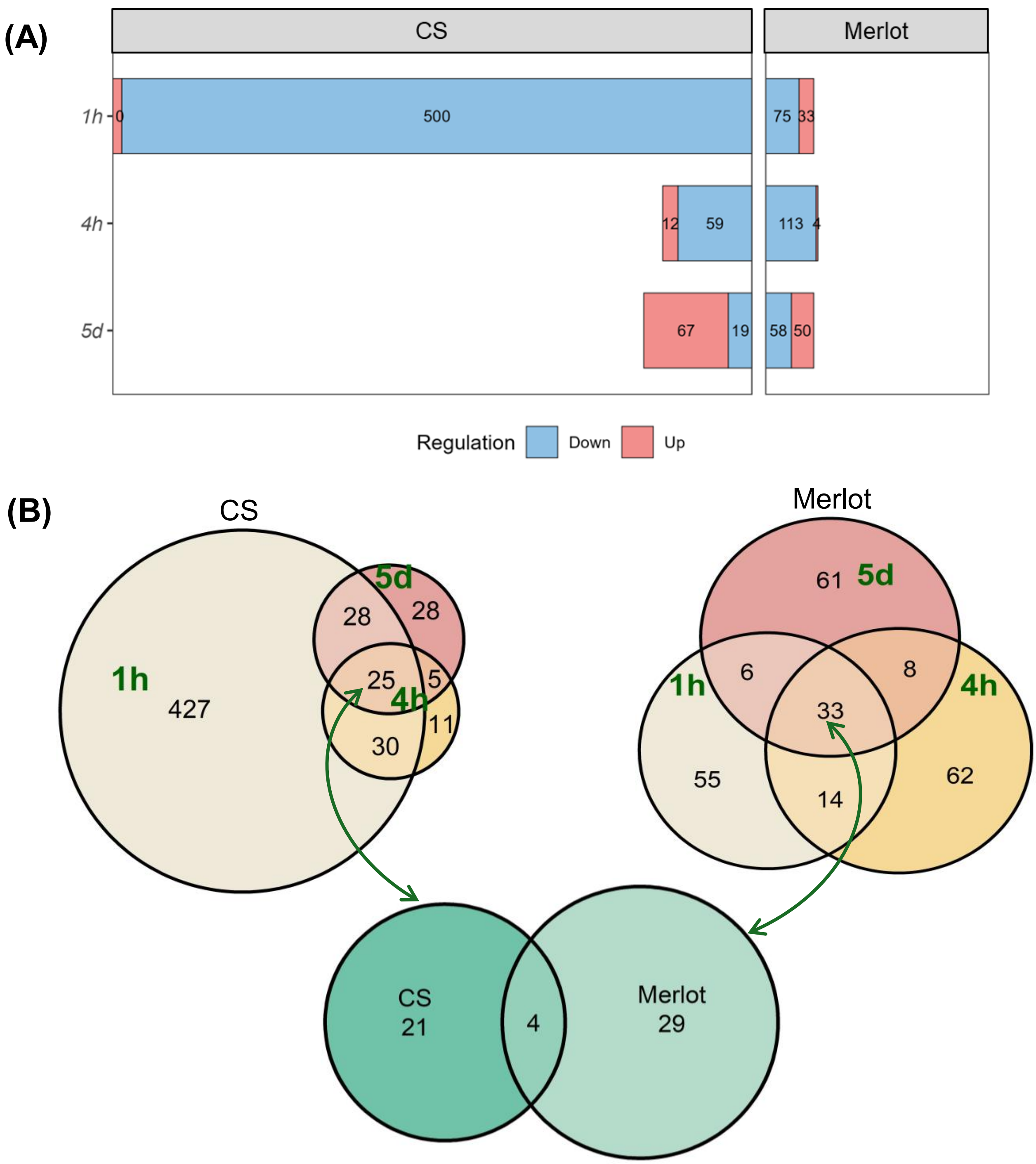


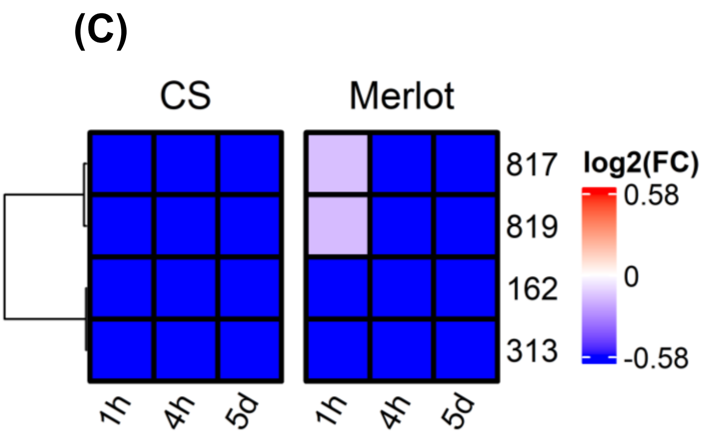


**Figure S3. (A)** Number of significant fragments identified by ***t*-test** at each time point ($p < 0.05$, raw) in Cabernet-Sauvignon (CS) and Merlot with untargeted GC-MS dataset. **(B)** Venn diagram showing the overlap of significant fragments across the three time points in both cultivars. **(C)** Heatmap illustrating the response patterns of overlapping fragments in CS and Merlot. Red indicates fragments increased by heat treatment, whereas blue indicates fragments decreased by heat treatment.

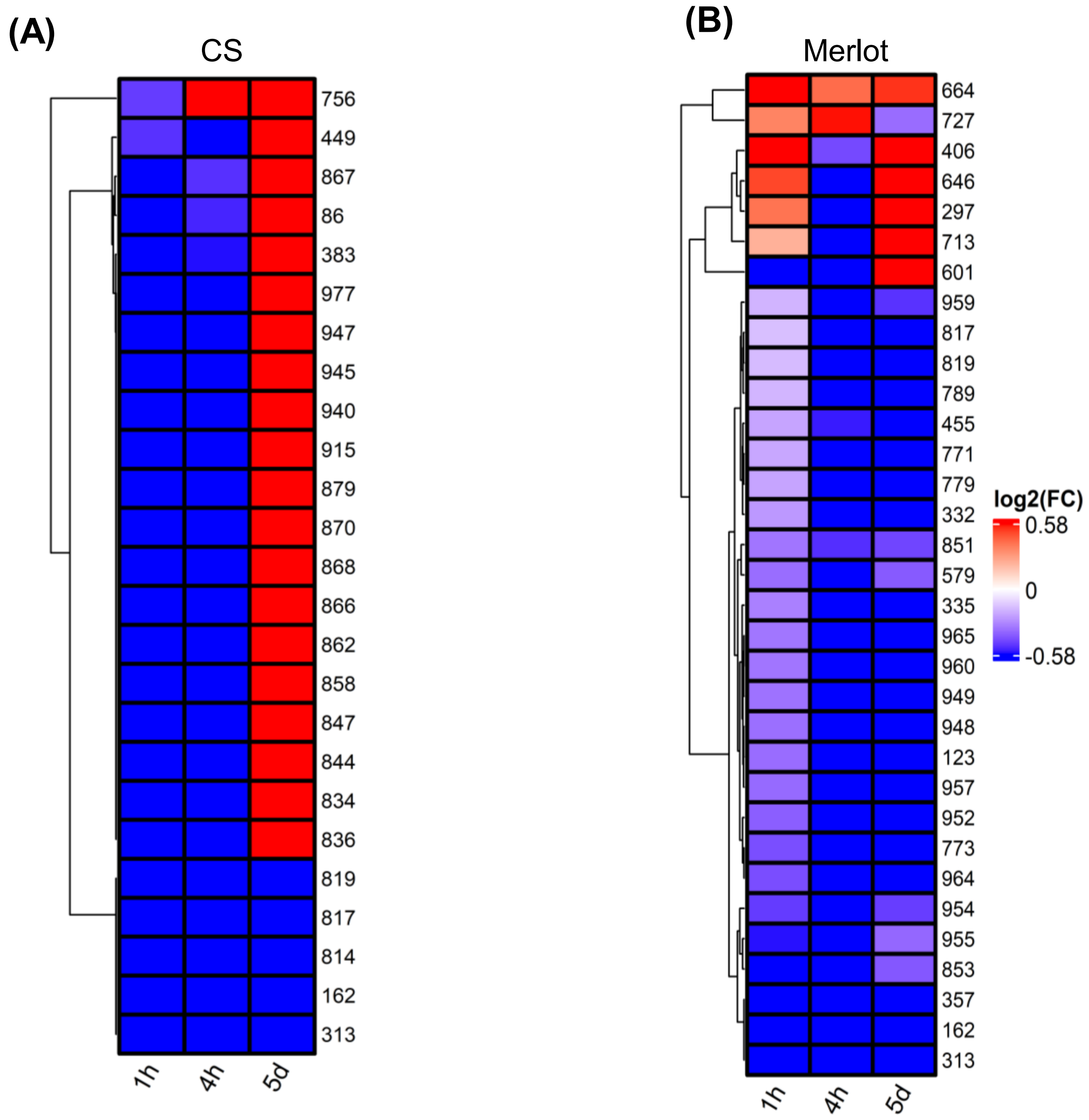


**Figure S4.** Heatmap illustrating the response patterns of overlapping fragments in Cabernet Sauvignon (CS) **(A)** and Merlot **(B)** across all time points. Red indicates fragments increased by heat treatment , whereas blue indicates fragments decreased by heat treatment.

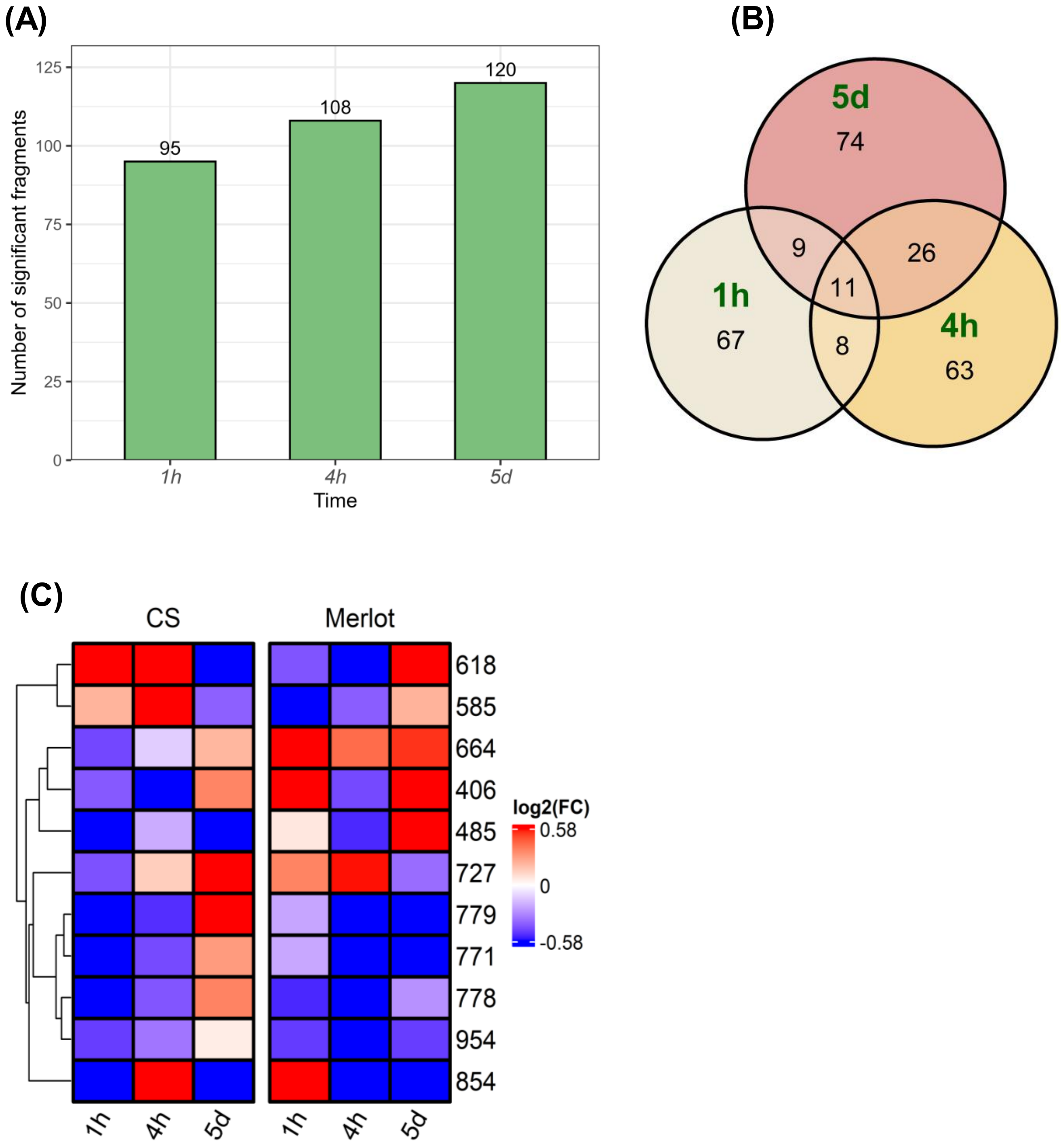


**Figure S5.** (**A)** Number of significant fragments identified by **two-way ANOVA** (Cultivar × Treatment interaction) at each time point (**p < 0.05, raw**). **(B)** Venn diagram showing the overlap of significant fragments across the three time points. **(C)** Heatmap illustrating the response patterns of overlapping fragments in Cabernet-Sauvignon (CS) and Merlot. Red indicates fragments increased by heat treatment (ht), while blue indicates fragments decreased by ht.
